# When Meaning Matters Most: Rethinking Cloze Probability in N400 Research

**DOI:** 10.1101/2025.04.29.651301

**Authors:** Yana Arkhipova, Alessandro Lopopolo, Shravan Vasishth, Milena Rabovsky

**Affiliations:** University of Potsdam, Germany

## Abstract

The N400 component of ERPs is modulated by how predictable a word is, but predictability is usually quantified with *lexical cloze*—the probability that readers supply that exact word in offline sentence completion tasks. This form-based metric is at odds with decades of evidence that the N400 is primarily sensitive to *meaning*. Here, we asked whether a measure of semantic feature predictability can better account for N400 amplitude modulation. We reanalysed two independent EEG datasets (N = 26 and N = 334), computing lexical and semantic cloze for each critical word. Across both datasets, semantic cloze emerged as a better predictor of the N400 data than lexical cloze. Using the same materials, we then compared semantic and lexical cloze with probabilities from four large language models (GPT-2, GPT-2.7b, RoBERTa, ALBERT). None of the LLM-derived predictors outperformed semantic cloze. Our findings support the view that the N400 primarily reflects semantic—not exact-word—processing. Methodologically, we argue that replacing lexical cloze with semantic cloze can substantially increase the explanatory power of N400 studies, and caution against substituting human norms with raw LLM probabilities.

## 1 Introduction

During language comprehension, humans rapidly form predictions about upcoming input based on contextual information (Altmann and Kamide, 1999). This predictive processing is fast, automatic, and serves as a crucial mechanism for facilitating comprehension. The influence of word predictability on overt processing behaviour has been observed across different modalities of language comprehension, including reading (Ehrlich and Rayner, 1981), listening (Altmann and Kamide, 2007; Kurumada et al., 2014), visual (Lieberman et al., 2018, in sign language comprehension), and haptic (Wilson et al., 2024, in braille reading). Given the ubiquity of prediction in human communication, any comprehensive theory of language comprehension must account for how people generate and update these anticipatory representations.

One of the most influential tools for investigating predictive processing is the N400 component of event-related potentials (ERPs). The N400 is a negative-going brain potential that peaks approximately 400 milliseconds after a stimulus, such as a word, is presented, and is elicited by every word in a sentence. Its amplitude is proportional to a word’s predictability: expected words elicit smaller N400 responses, whereas unexpected words produce larger, more negative amplitudes. This sensitivity to expectancy, together with the high temporal resolution of ERP methods that allow researchers to precisely track predictability effects at the scalp, has made the N400 a critical tool for investigating how humans process language in real-time. Insights gained from N400 research have shaped many contemporary models of language comprehension (e.g. Rabovsky et al., 2018; Brouwer et al., 2017; Fitz and Chang, 2019; Cheyette and Plaut, 2017; Laszlo and Plaut, 2012) and continue to inform the field.

Given the pivotal role that the N400 plays in language comprehension research, it is crucial that the measurements we use to study this component are as precise and reliable as possible. Cloze probability (Taylor, 1953), which estimates how likely a particular word is in a given sentence, has long been considered the primary predictor of N400 amplitude. While this correlation holds true to a significant extent, there are well-documented exceptions. We will argue that, based on current empirical findings about the N400, cloze probability is not an optimal choice of predictability measurement in N400 studies, and should be replaced by a semantic feature predictability measurement.

To that end, we propose a semantic feature predictability measure and compare it directly to lexical cloze. Additionally, given the recent developments in using large language models (LLMs) as alternatives to human-derived norms for predicting N400 amplitude (e.g., Heilbron et al., 2022; Michaelov et al., 2022), in the second part of this study we evaluate our semantic measure against LLM-generated cloze.

In the next section, we first review how word predictability has traditionally been operationalized through cloze probability and examine whether this measure can adequately account for the N400 amplitude change. We then introduce and test an alternative measure—“semantic cloze”—that aligns more closely with contemporary understanding of the N400.

### Cloze Probability

Word predictability is operationalized as its cloze probability. It is typically obtained through a cloze task, where participants read truncated sentences and supply the first word that comes to mind (Taylor, 1953).The cloze probability of a target word is then the proportion of responses that match that target word exactly. Consequently, cloze probability is inherently ‘lexical’—that is, it reflects the probability of encountering that specific word in the given context.

Because it serves as direct index of word predictability, cloze probability plays a pivotal role in the design of most N400 experiments. Researchers often manipulate the factor “congruence” by selecting stimuli with either high (‘congruent’) or low (‘incongruent’) cloze probability target words, discarding or retaining items based on cloze test results. Other experimental paradigms group stimuli into graded-expectancy conditions (e.g., high, medium, and low cloze) entirely on the basis of cloze scores. Consequently, cloze probability not only shapes the stimuli but often also determines how the experimental conditions themselves are defined.

At the analysis stage, cloze probability often functions as a predictor variable in mixed-effects models, directly influencing model fitting and the estimation of the effect of interest. Even when it’s not used as a continuous predictor, its impact persists through prior decisions about stimulus selection and condition assignment, ultimately carrying over into the statistical analysis via binary predictors.

Given its importance at every step—experimental design, item selection, and statistical analysis—predictability should ideally be measured with as much precision as possible. Imprecise measurement can lead to spurious results or contribute to inconsistencies across N400 studies.

### Semantic and Lexical Predictability in ERPs

Graded attenuation of the N400 as a function of cloze probability is arguably the most well-known and widely replicated effect in language ERP research (e.g Federmeier and Kutas, 1999; Frade et al., 2022, and many others). The observation that the N400 becomes more negative as word predictability decreases is often regarded as the “ground truth” for this component. However, it is also well-documented that the N400 may still be attenuated for items with 0% cloze probability.

Studies that use the semantic relatedness manipulation have consistently found that the N400 can be attenuated to words with very low cloze probabilities (Federmeier and Kutas, 1999; Arbel et al., 2011; Thornhill and Van Petten, 2012; Kwon et al., 2017). In these studies, participants read sentences designed to elicit specific expectations, e.g. *‘When you inhale, air enters your…’*, which primes the reader to predict *‘lungs’* (Arbel et al., 2011). The sentence then ends in either highly expected, high-cloze continuation—*‘lungs’*; an unexpected ending with 0% cloze probability—*‘money’*; and an unexpected, but semantically-related continuation— *‘heart’*. If cloze probability alone predicted N400 amplitude, *‘money’* and *‘heart’* would have similarly large N400s. What has been found, however, is that the N400 shows attenuation in response to *‘heart’*, likely because it shares some semantic features with *‘lungs’* (both being vital organs). The standard cloze probability measurement would fail to capture this effect.

This semantic facilitation effect suggests that the N400 is more accurately understood as a correlate of semantic, rather than exact-word, predictability. On this view, the sentence *‘When you inhale, air enters your…’* would trigger preactivation of upcoming semantic features (e.g. [+organ], [+breathing]). The word *‘lungs’* would then receive the most amount of attenuation in the N400 window as it matches both features. The word *‘heart’* would receive only an intermediate amount of attenuation since it only partially matches those features ([+organ],[-breathing]).

The N400’s sensitivity to semantics is further corroborated by research involving bilingual populations. For example, Blackburn and Wicha (2022) asked Spanish-English bilinguals to read Spanish sentences containing a predictability manipulation. The target words, either expected or unexpected, appeared either in Spanish or in English. Thus, the target words shared only semantic meaning but not lexical overlap. The results showed that the N400 attenuated equally for both Spanish and English endings. Similar results have been reported in other bilingual studies (Moreno et al., 2002; Valdés Kroff et al., 2020). This again demonstrates that semantic overlap, not lexical overlap, drives N400 attenuation.

This characterization of the N400 as a marker of semantic pre-activation is also reflected in current models of language comprehension. For instance, the Sentence Gestalt model (Rabovsky et al., 2018) views the N400 as the index of change in a representation of sentence meaning, and explicitly links reduced N400 effects to partial semantic feature overlap. The Retrieval-Integration account (Brouwer et al., 2017) proposes that the N400 reflects the retrieval of word meaning. While the two models propose different mechanisms behind the N400 modulation, they converge on attributing the component to meaning processing.

Nevertheless, some evidence for exact-word prediction in the N400 window has been reported, though the findings are less consistent. For example, studies using the pre-nominal mismatch paradigm are often cited as strong evidence for the pre-activation of specific lexical items. In DeLong et al. (2005), participants read sentences like *‘It was a windy day, so the boy went out to fly a/an kite’* while ERPs were recorded at the pre-nominal article (a/an). The article either matched or mismatched the most likely upcoming noun (*‘kite’*). The ERPs showed an increased negativity in the N400 window in response to the mismatching article, which was taken to suggest that readers must have pre-activated the specific lexical item *‘kite’*, including its phonological form, to know whether *‘a’* or *‘an’* would be phonologically appropriate. These results are not only taken to support the idea that comprehenders predict information at all levels of representation (meaning, word form, phonology; what is known as ‘strong prediction view’), but that the consequences of violation of those predictions are reflected in the N400 specifically.

While the DeLong and colleagues’ study could indeed be taken to offer strong evidence for word-form pre-activation in the N400 window, this effect appears to be somewhat unreliable. Several studies to date have failed to replicate the pre-nominal pre-activation effect (Nieuwland et al., 2018; Kochari and Flecken, 2018), or yielded inconclusive results (Nicenboim et al., 2020). Others have reported ERP effects at different latencies and topographical distributions than the N400 (Nieuwland et al., 2020). Even if these effects were confined to the N400, they do not have to necessitate word-form pre-activation, as demonstrated via simulations with the SG model Rabovsky (2020) and discussed in Nicenboim et al. (2020), and could instead index change in a probabilistic representation of meaning triggered by the mismatching article.

Still, effects of lexical predictability in the N400 window cannot be entirely ruled out. For example, Ito et al. (2016) demonstrated that some word-form pre-activation may be reflected in that time window, but only when the sentence is sufficiently constraining. In their study, words that were physically similar to highly predictable words (e.g., *‘book’* and *‘hook’*) elicited attenuated N400 responses relative to unrelated controls (e.g., *‘sofa’*). Importantly, this effect occurred only for *≈* 100% cloze probability target words, and no such attenuation was found for medium-predictability items.

In sum, the experimental evidence and modeling implementation of the N400 reviewed so far suggests that the current operationalization of cloze probability as a measure of exact-word prediction is not an optimal predictor of the N400.

As can be seen, the experimental evidence—and the prevailing understanding of the N400—are at odds with the exact-word prediction implied by cloze probability measurements. While cloze probability reflects the predictability of a specific word, the N400 appears to be primarily sensitive to the preactivation of semantic features. Of course, exact-word predictability may often coincide with predicted semantic features (e.g., in 100% cloze probability cases where participants unanimously choose the same word). However, this alignment is not guaranteed, particularly for medium-cloze items. For instance, one can easily imagine half of the cloze task participants completing the sentence *‘After a long day at work, she collapsed onto the comfortable…’* with *‘couch’*, while the other half provide *‘sofa’*. Using the current operationalization, cloze probability for *‘couch’* alone would then be 50%, classifying it as a medium-cloze item. Yet both *‘couch’* and *‘sofa’* share the semantic feature ‘long, comfortable seat’, making that concept effectively 100% predictable despite the split lexical choices. Consequently, in such cases, standard cloze probability measures may underestimate semantic predictability.

## 2 The Current Study

This study investigates whether the N400 amplitude change can be better accounted for by predictability of semantic features as compared to lexical predictability.

First, we propose and evaluate a new measurement of cloze probability—what we term ‘semantic cloze’. To this end, we reanalyse ERP data from two datasets: Arkhipova et al. (2019) and Nieuwland et al. (2018), using the original cloze-task responses to calculate both semantic and lexical cloze values for each target word (see below for details). We then compare how well the two measures account for N400 amplitude. Our results show that semantic cloze provides a superior fit relative to conventional (lexical) cloze probability.

In the second part of this work, we further test this finding by evaluating whether cloze probability generated by large language models (LLMs) can provide a superior fit to the N400 data than semantic cloze. Given claims that LLMs capture human-like linguistic patterns (Goldstein et al., 2022; Michaelov et al., 2022) it remains an open question whether their probability estimates provide a more accurate reflection of real-time language processing than traditional cloze measures. To test this, we compare model-derived probabilities with human-generated semantic and lexical cloze estimates. We find that LLM-derived predictions do not outperform semantic cloze in accounting for N400 amplitude.

### 2.1 Method

### 2.2 Overview of Existing Data

We reanalyzed two existing datasets from independent ERP experiments: Arkhipova et al. (2019)—henceforth ARK, and (Nieuwland et al., 2018)—henceforth NI18. These datasets were chosen because each contained both EEG recordings and raw cloze data, which were necessary for estimating semantic cloze.

In both studies, native English speakers read sentences presented one word at a time in the center of a screen, while EEG was recorded. Additional details regarding participant demographics, materials, and procedures are summarized in Table 1.

**Table 1.** Summary of cloze task and lexical-semantic analysis results. Number of cloze task participants is displayed in ranges as the number of participants varied across lists of stimuli (ARK had four counterbalanced lists)

| Data | N Subj |  | Critical words | Cloze type | Mean (SD) | Difference Sem-Lex |  | Sem Cloze Interrater reliability (ICC), 95% CI |
| --- | --- | --- | --- | --- | --- | --- | --- | --- |
|  | Cloze Task | EEG |  |  |  | Mean(SD) | Range |  |
| ARK | 22-27 | 26 | verbs<br>adjectives | Lex | 0.21 (0.24) | 0.14 (0.18) | 0–0.78 | ‘Long’ Condition 0.994 [0.992, 0.996] |
|  |  |  |  | Sem | 0.35 (0.33) |  |  | ‘Short’ Condition 0.908 [0.834, 0.943] |
| NI18 | 29-30 | 334 | nouns | Lex | 0.48 (0.41) | 0.07 (0.14) | 0–0.78 | 0.962 [0.95, 0.97], $p < 0.001$ |
|  |  |  |  | Sem | 0.55 (0.41) |  |  |  |

The experimental manipulations in the original studies are not important for the present analysis, as we are interested in examining the N400 change as a function of lexical or semantic predictability. Therefore, we conduct statistical analyses on all target words in these datasets—regardless of whether they were predictable or unpredictable. However, in some of the plots that follow, data are displayed by condition. We therefore briefly clarify the relevant manipulations:

**ARK Dataset**: This study examined how quickly comprehenders can reverse their expectations when encountering the word ‘nonetheless’ in a sentence. Thus, in addition to predictability manipulations, this dataset contained a ‘Distance’ manipulation. This refers to whether the target word appeared immediately after ‘Nonetheless’ (e.g. *He had a map with him. He was, <u>nonetheless, lost</u>*—‘Short’ condition) or with intervening words in between (e.g. *<u>Nonetheless</u>, he was <u>lost</u>*—‘Long’ condition), where the ‘Short’ condition would require the participant to reverse the expectations immediately to anticipate the verb. Example set of stimuli can be found in the Supplementary Materials (Table S1).

The **NI18 Dataset** was a replication study examining ERP effects at prenominal articles (a/an). Apart from the article expectancy manipulation, the predictability of the noun also varied by condition. Here, we focus only on the noun, as it carries semantic content (unlike the indefinite articles a/an). Table S2 in the Supplementary Materials provides example sentences illustrating this design.

### 2.3 Semantic and Lexical Cloze Calculation

Cloze task responses for both ARK and NI18 were obtained by the respective authors through a standard cloze task procedure. Sentences were truncated at the critical word and distributed across four lists using a Latin square design. Participants were asked to complete each sentence with the most suitable continuation.

Lexical cloze was calculated as the proportion of responses that exactly matched the critical word presented in the ERP experiment, following the standard cloze probability estimation proposed by Taylor (1953). Note that our lexical cloze calculation for one sentence (#79) in NI18 differs from that provided by the original authors. That item was divided by 30 although a participant left the response blank. We recalculated that proportion over 29 responders. Other changes are described in their respective sections, and are additionally summarised in Table S5.

Semantic cloze was calculated as the proportion of responses that were semantically similar to the critical word, given the sentence context.

The calculation logic is illustrated in Fig.1. In this example, 6 out of 25 participants completed the sentence with ‘persisted’, which was the exact-word match to the critical word presented in the EEG experiment, giving it a 24% lexical cloze probability value. For semantic cloze, all semantically similar continuations (all words within the blue dashed rectangle in Fig.1), including ‘persisted’, were counted, resulting in a 56% semantic cloze probability for the same critical word.

**Figure 1.**
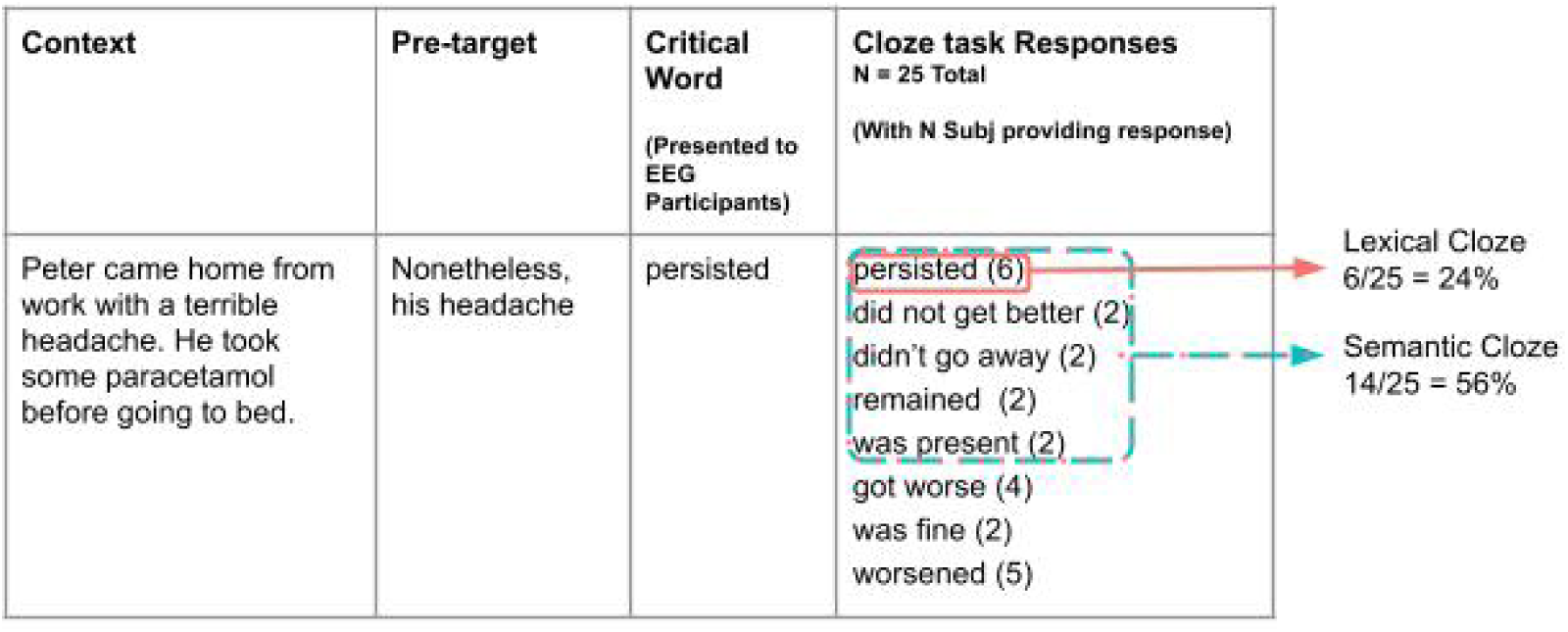
Illustration of lexical and semantic cloze calculation logic. Rightmost column shows responses provided by the cloze task participants, with the number of participants that provided each response in brackets. All instances of ‘persisted’ (6 total) count towards lexical cloze. Words in the blue dashed rectangle count towards semantic cloze for this target word (14 total).

Each target word, therefore, had associated lexical and semantic cloze values derived from the same set of responses. By definition, semantic cloze is either equal to or greater than the lexical cloze value.

To ensure the reliability of semantic cloze measurement, two raters independently evaluated the raw cloze responses for each sentence and identified all responses that constituted a close semantic match to the critical word in the given context. Each rater calculated the semantic cloze probability by dividing the number of semantically similar responses by the total number of responses per item. Interrater agreement was excellent, as indicated by the intraclass correlation coefficient (see Table 1). (Note that the second rater was different for each condition in ARK, which is why ICCs are reported separately for the “Long” and “Short” conditions.) The mean semantic cloze value from the two raters was then computed for each item and used in all subsequent analyses.

Descriptive statistics for lexical and semantic cloze values are summarised in Table 1. Semantic and lexical cloze values were highly correlated (ARK: *r* = .86; NI18: *r* = .94), with semantic cloze invariably equal to or greater than lexical cloze. Figure 2 illustrates the distribution and differences between the two measures. In the NI18 dataset, the two measures were more similar (mean difference = .07, SD = .14, range = 0–.78), likely because many sentences primed concrete nouns (e.g., ‘The old wives tale says that if you want to keep the doctor away then you should eat…’). Such contexts offer fewer opportunities for semantically related alternatives, causing participants to supply the exact word (‘apple’) and minimizing the gap between semantic and lexical cloze. By contrast, the ARK dataset included verbs and adjectives as critical words, which permitted more varied completions and thus led to a greater discrepancy between the two cloze measures (mean difference = 0.14, SD = 0.18, range = 0-.78).

**Figure 2.**
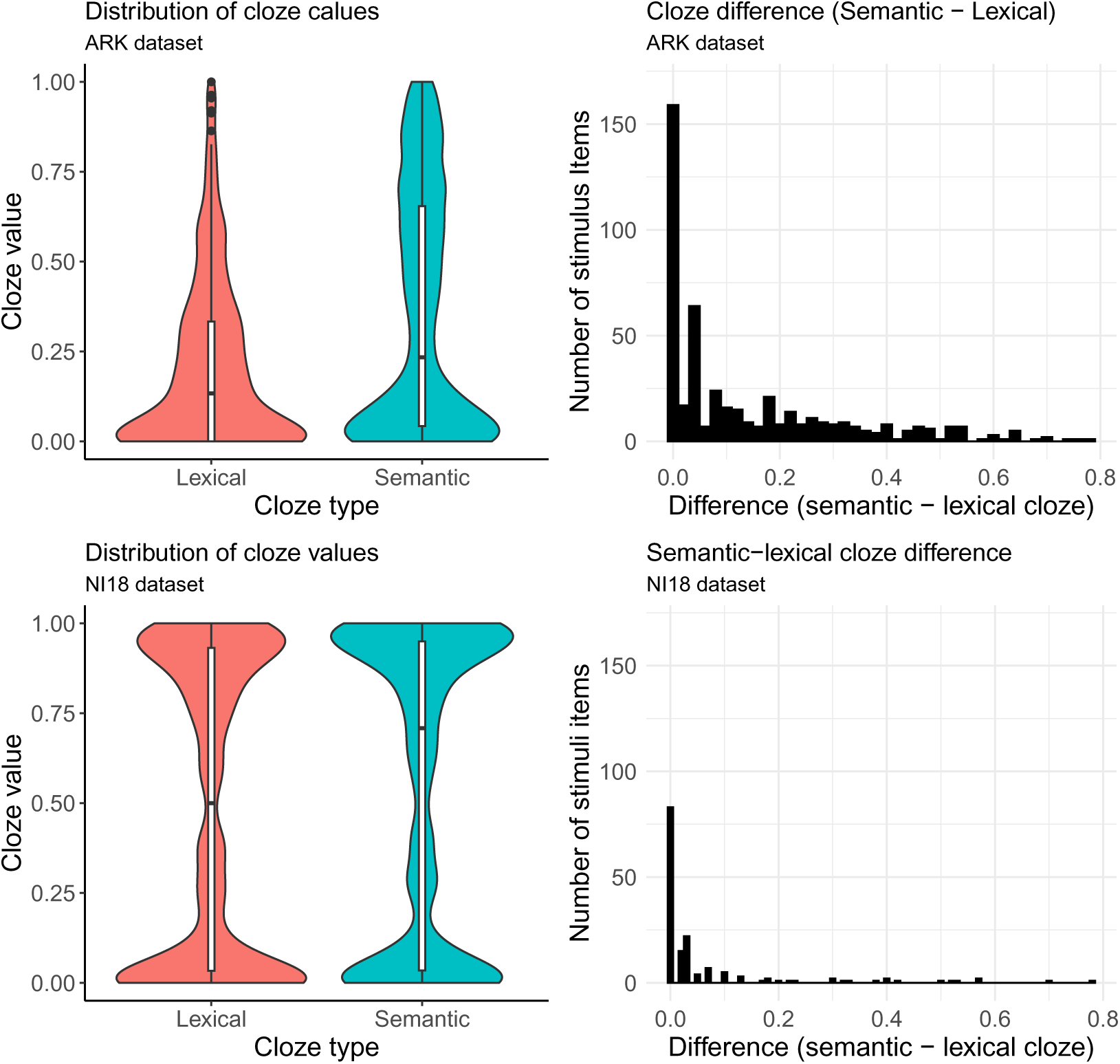
**Left:** Distribution of lexical and semantic cloze probabilities with superimposed box plots. **Right:** Distribution of differences between semantic and lexical clozes, indicating that semantic cloze probabilities are often higher than lexical cloze probabilities.

### 2.4 EEG Pre-processing

#### ARK Dataset

EEG data were pre-processed in Matlab (2021a) using EEGLAB and ERPLAB toolboxes. Continuous data were downsampled to 1000 Hz using a polyphase antialiasing filter, and bandpass filtered using a second-order Butterworth infinite impulse response (IIR) acausal filter (0.1–30 Hz). Next, the data were re-referenced to the average of the left and right mastoids and epoched into segments spanning −200 to 1000 ms around the onset of the critical word. The epochs were baseline corrected relative to the 200 pre-stimulus time window.

Automatic artifact rejection was performed with EEGLAB’s eegthresh() function, using a −80/+80 *µ*V threshold within the −200 to 800 ms time window. To preserve as many segments as possible, trials were rejected only if artifacts were detected in centroparietal or occipital channels (Cz, Pz, Oz, C3, C4, CP5, CP6, CP1, CP2, P7, P8, P3, P4, PO3, PO4, O1, O2), since frontal electrodes were not of primary interest. Of the total 3120 trials, 53 (1.7%) contained artifacts and were excluded from analysis.

#### NI18 Dataset

We used the pre-processed and segmented data provided by the original authors (available at https://osf.io/eyzaq/files/osfstorage) to extract N400 trial means, introducing only minor changes from the original NI18 to match the settings in the ARK dataset. Specifically, we applied a −200 ms baseline correction (instead of −100 ms). In NI18, the −100 ms baseline was selected to match the replicated study’s design; however, a −200 to 0 ms baseline is more common in N400 research.

When exporting the data after baseline correction, we found that two trials in the dataset contained mis-timed trigger codes (typically caused by a hardware glitch); we excluded those two trials, as it was not clear whether the triggers marked the word onset correctly. Consequently, the final data set contains 25 846 data points (vs. 25 848 in the public NI18 data). This did not affect our results (see S3 and S4 in the Supplementary Material.)

### 2.5 Statistical Analyses

All analyses were performed in R (v4.3.3; R Core Team, 2023) using the lme4 package (Bates et al., 2015) for linear mixed-effects modeling and bbmle (Bolker and Team, 2023) for AIC comparisons.

Our goal was to determine whether the predictability of semantic features or lexical items better explains N400 amplitude modulations. To this end, we conducted two analyses on each dataset.

#### Joint Model vs. Reduced Models

First, we assessed whether each of the cloze measures contributes unique explanatory power to the N400. We specified a joint model that included both lexical and semantic cloze as predictors. We attempted to fit “maximal” models, following the recommendations of Barr et al. (2014). For ARK: with fixed effects for semantic and lexical clozes and distance, and random intercepts for and slopes for the joint effect of semantic and lexical clozes. For NI18, the model was the same except for the factor Lab included instead of Distance. After these models failed to converge, we systematically simplified the random-effects structure by removing random slopes for items and then for participants. The final model structure included random intercepts for participants and items:

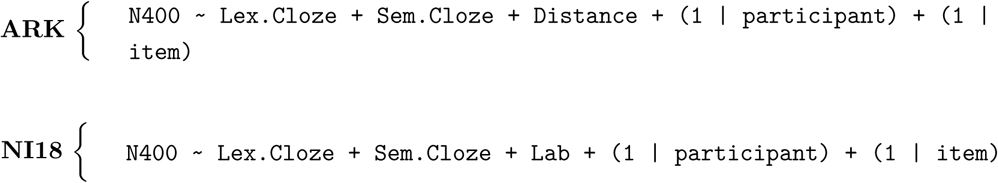

To check for potential collinearity, we computed variance inflation factors (VIFs) for each model. All VIFs were less than 4, which is well below the conventional threshold of 10 (Cohen et al., 2013). We then compared each joint model to two reduced models, omitting lexical or semantic cloze, respectively, via likelihood ratio tests.

#### AIC Comparisons

Our second analysis compared model fits using the Akaike Information Criterion (AIC). AIC quantifies information loss in a given model and enables evaluation of predictive accuracy for non-nested models (Burnham et al., 2011). Lower AIC values indicate better fit. The difference (Δ*_i_*) between the lowest AIC model and any other candidate model serves as a measure of relative support. Models with Δ*_i_ >*10 have essentially no support, Δ*_i_* = 4 to 7 – considerably less support, and Δ*_i_* = 0 to 2 – substantial support.

Because the ARK dataset had a relatively small sample size, we used AIC corrected for small samples (AICc; Hurvich and Tsai, 1989), whereas we used regular AIC for the NI18 dataset.

Additionally, we report fixed-effect estimates from the linear mixed-effects models. To evaluate significance, each model was compared to a reduced model (without the fixed effect in question) using the anova() function in R, which performs a likelihood ratio test. We used an alpha level of 0.05 for all statistical tests.

#### Variables and Transformations

For both datasets, the N400 measure was the mean amplitude in the 300–500 ms window after stimulus onset, averaged over a posterior electrode cluster (CP1/2, C3/4, Cz, Pz, P3/4)^1^, where the N400 effect is typically maximal (Kutas and Federmeier, 2011). Cloze predictor variables were z-scored and centred. No transformations were applied to the N400 mean amplitude data.

All reported *β*-coefficients and 95% confidence intervals (CIs) were transformed back from the z-scored scale to raw values, representing the voltage change associated with moving from 0% to 100% cloze probability. This back-transformation was achieved by dividing the standardized coefficient by the standard deviation of the original (non-standardized) predictor, yielding interpretable estimates in microvolts. CIs were computed using the Wald method.

### 2.6 Results

#### ARK Dataset

##### Joint Model

A log likelihood ratio test comparing the joint model to the model containing only semantic cloze showed that adding lexical cloze to a model already containing semantic cloze did not significantly improve the fit, *χ*^2^(1) = 1.26, *p* = 0.261. In contrast, the joint model showed a significant improvement over the lexical-only model, *χ*^2^(1) = 4.9, *p* = 0.027, suggesting that the model including semantic cloze as well as lexical cloze furnishes a better fit compared to the model including only lexical cloze.

##### LMEM

For the ARK dataset, lexical cloze had no significant effect on N400 amplitude (*β* = 1.94, 95% CI = [-0.55, 4.4], *χ*^2^(1) = 2.3, *p* = 0.1). In contrast, semantic cloze showed a statistically significant effect (*β* = 2.2, 95% CI = [0.43, 3.9], *χ*^2^(1) = 5.9, *p* = 0.01). The positive *β* coefficient suggests that higher semantic cloze probabilities are associated with less negative N400 amplitudes.

##### AICc Model Comparison

The AICc values and weights were computed using the ICtab() function of the bbmle package, specifying the type as ‘AICc’. The results are summarised in Table 2.

**Table 2.** AICc comparison for the ARK dataset.

| | AICc | ΔAICc | df | weight |
| --- | --- | --- | --- | --- |
| Semantic Cloze model | 26205.91 | 0.0 | 6 | 0.86 |
| Lexical Cloze model | 26209.54 | 3.6 | 6 | 0.14 |

The AICc analysis indicates that the semantic cloze model provides the best fit for the N400 data, as reflected by its lower AICc value. The lexical cloze model’s ΔAICc of 3.6 suggests considerably less support relative to the semantic cloze model. Moreover, the model weights support this conclusion, with semantic cloze carrying a weight of 0.86—implying an 86% probability of being the superior model among those compared (Burnham & Anderson, 2001).

##### Grand Average ERPs

Figures 3 and 4 display grand-average ERPs of the same dataset, binned according to either lexical or semantic cloze values. First, we sorted the stimuli in ascending order based on lexical cloze and divided them into four equal-sized groups—Low, Low-Medium, Medium-High, and High—each containing 30 sentences. Next, we re-sorted these same stimuli in ascending order by their semantic cloze values and again partitioned them into four groups. We then plotted separate grand-average

**Figure 3.**
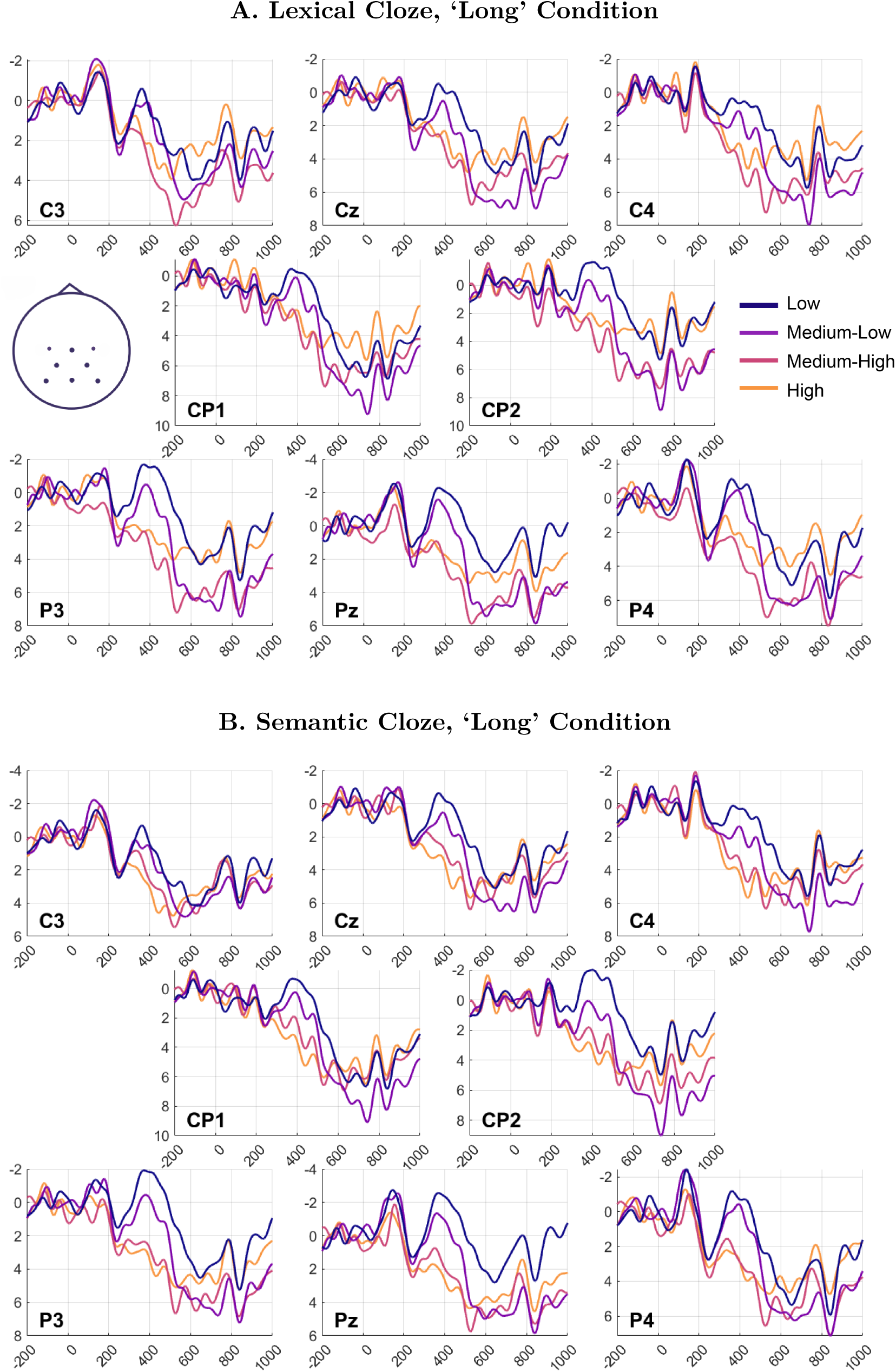
Grand average ERPs by each ROI channel, ‘Long’ Condition: Lexical Cloze (A); Semantic Cloze (B). ERPs were 15 Hz filtered for presentation.

**Figure 4.**
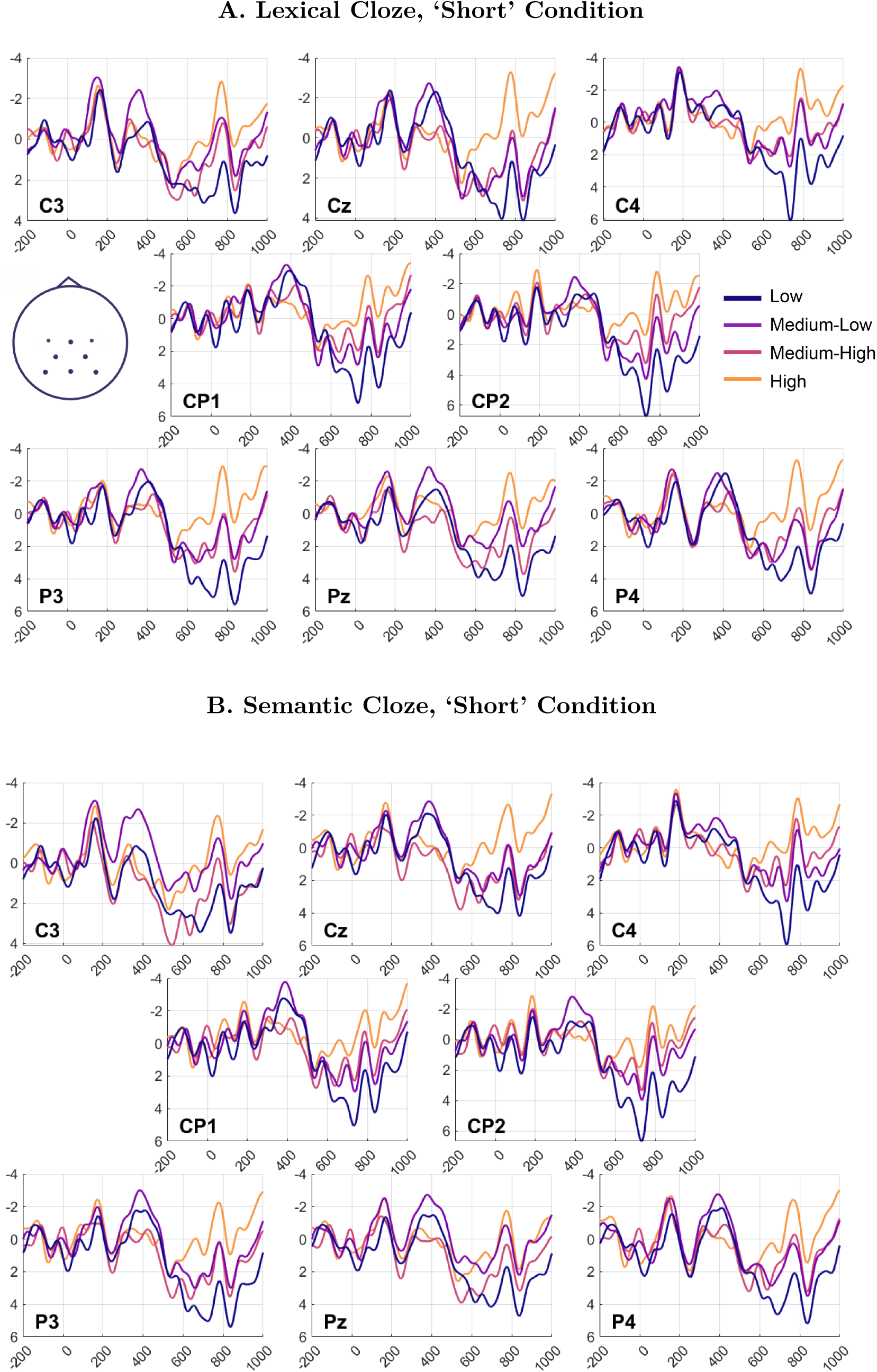
Grand average ERPs by each ROI channel, ‘Short’ Condition: Lexical Cloze (A); Semantic Cloze (B). ERPs were 15 Hz filtered for presentation.

ERP waveforms for each sorting scheme, resulting in distinct ERP plots for the lexical versus semantic bins.

Figures 6 and 5 display mean N400 amplitudes binned by lexical and semantic cloze, separately for the Long and Short conditions. While we are not interested in the effect of condition, plotting them separately was necessary to address baseline differences. Bar heights represent the average N400 amplitude for each bin, and error bars indicate the standard error of the mean.

**Figure 5.**
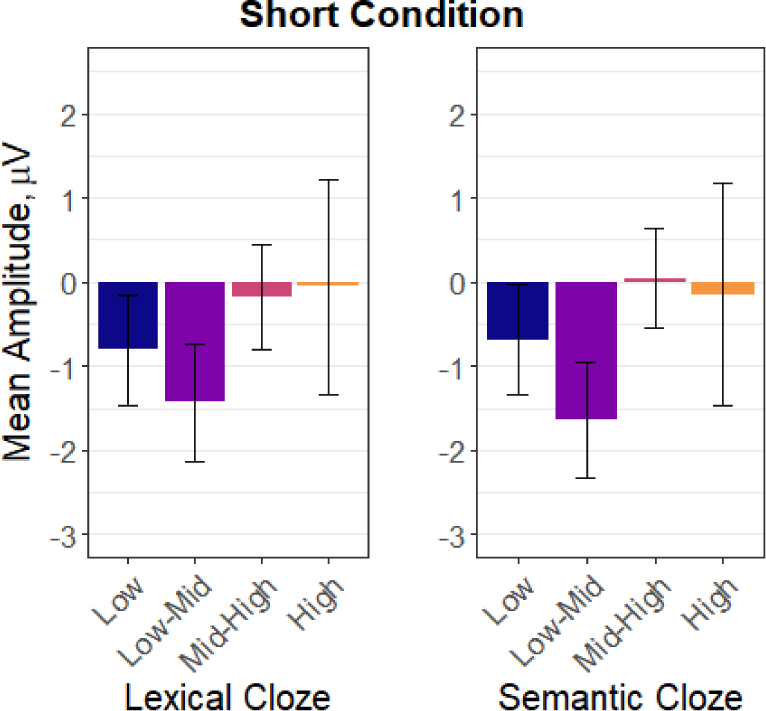
N400 means and SE in the Short distance condition.

**Figure 6.**
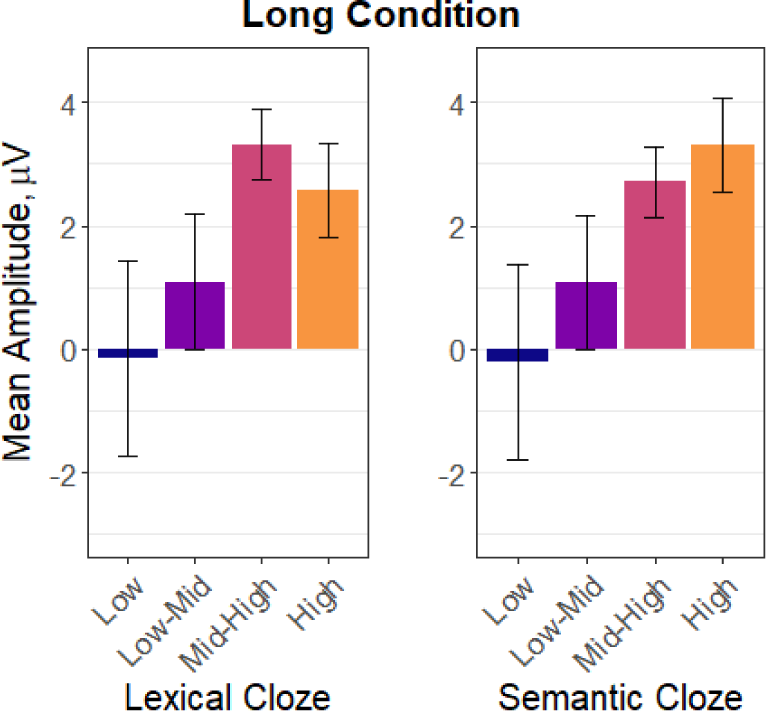
N400 means and SE in the Long distance condition.

From both the grand-average ERPs and the bar graphs summarising the long condition, the largest effect of semantic cloze appears in the High and Mid-High bins, where it produces a more clearly graded pattern of N400 attenuation than lexical cloze. When binned by lexical cloze, the highest predictability group appears to have lower amplitude than the mid-high predictability items. This pattern visually supports our finding that semantic cloze is a stronger predictor of N400 amplitude than lexical cloze, aligning with the broader conclusion that comprehenders primarily pre-activate upcoming semantic features rather than fully specified words. However, this improved graded pattern does not appear to hold in the short condition. Consequently, we tested for a potential interaction between cloze type and Distance condition to further clarify these observations.

##### Distance x Semantic Cloze

We compared a model with only the main effects of semantic cloze and distance to one including their interaction. The interaction model fit the data slightly better (*χ*^2^(1) = 3.27, *p* = 0.07), further reducing the AICc by 2.1 (AICc = 26203.8) compared to the no-interaction model (AICc = 26205.9). The interaction, however, did not reach the conventional alpha level (*p* = 0.05).

We used the marginal trends analysis (via the emmeans package to examine the effect of semantic cloze in each distance condition. For the long condition, semantic cloze remained a significant predictor of the N400 (*β* = 1.28, SE = 0.43, 95% CI = [0.44, 2.11]). However, in the Short condition, the slope of the semantic cloze was not significant (*β* = 0.19, SE = 0.42, 95% CI = [-0.63,1.02]), as evidenced by the confidence interval including zero.

### NI18 Dataset

#### Joint Model

To evaluate whether each predictor contributed unique explanatory power to the N400 amplitude, we compared a joint model with both semantic and lexical cloze to two reduced models that omitted either semantic or lexical cloze.

Comparing the joint model to a model containing only semantic cloze yielded a non-significant difference in log-likelihood *χ*^2^(1)= 0.0884, *p* = 0.77, indicating that adding lexical cloze to a model already containing semantic cloze did not significantly improve model fit.

In contrast, the difference in deviance between the joint model and the lexical-cloze-only model was significant, *χ*^2^(1)= 18.99, *p <* 0.001, suggesting that adding semantic cloze to a model already containing lexical cloze substantially improved the model’s fit to the data.

#### LMEM

For the NI18 dataset, both lexical cloze (*β* = 2.65, 95% CI = [2.35, 2.96], *χ*^2^(1) = 287.24, *p <* 0.001), and semantic cloze (*β* = 2.76, 95% CI = [2.45, 3.07], *χ*^2^(1)= 306.14, *p <* 0.001) significantly predicted N400 amplitude. In both cases, higher cloze values were associated with more positive N400 amplitudes.

#### AIC Model Comparison

We computed AIC values and weights using the ICtab() function from bbmle (with type = ‘AIC’). The results are summarized in Table 3. The ΔAIC of 18.8 indicates that the lexical cloze model has essentially no support relative to the semantic cloze model.

**Table 3.**
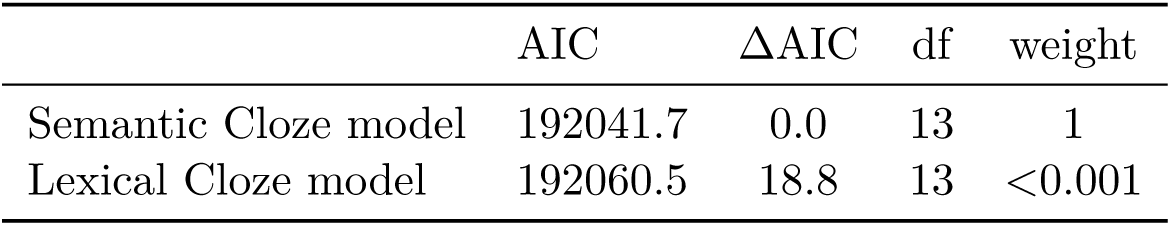
AIC comparison for the NI18 dataset.

### 2.7 Discussion

Our analyses revealed that semantic feature predictability, as captured by semantic cloze, emerged as a stronger predictor of N400 amplitude than lexical cloze. This result held for verbs and adjectives (in the ARK dataset) as well as for nouns (in the NI18 dataset), suggesting that comprehenders pre-activate upcoming semantic features for both abstract qualities and concrete objects, and that this pre-activation is captured in the N400 component. Notably, semantic cloze substantially improved model fit even in the NI18 dataset, where semantic and lexical cloze values differed only slightly.

Our additional, by-condition analysis in ARK indicated that semantic cloze was a significant predictor only in the Long condition, with no reliable effect in the Short condition. Although the interaction of cloze and distance was not significant, this pattern might suggest that predictability effects are reduced under time constraints, giving comprehenders limited time to “reverse” any initial expectations in the Short condition—consistent with suggestions that prediction requires sufficient time to develop (Chow et al., 2018; Liao and Lau, 2020). Given the small sample size in ARK, however, these findings should be interpreted cautiously. Even so, the overall superior AICc and AIC metrics across both ARK and NI18 support the conclusion that semantic cloze is a more effective predictor of N400 amplitude than lexical cloze.

However, one might wonder whether our findings simply reflect the limitations inherent to human-generated cloze responses, rather than a fundamental difference between lexical and semantic predictability. It has been argued that the small sample sizes typically used in cloze tasks (around 30–50 participants) may be inadequate for accurately estimating the predictability of unexpected (i.e., low-cloze) items (Szewczyk and Federmeier, 2022; Shain et al., 2024). In highly constraining contexts, participants tend to produce the most predictable words, even when multiple low-probability continuations are possible. The authors argue that it is impossible to improve precision in those estimates using the human-generated responses, as it would require a prohibitively large number of participants.

Large language models (LLMs) have been proposed as a remedy for such potential biases, given that they can generate probability distributions over a vast space of possible completions and may therefore be capable of obtaining more precise cloze values, particularly for low-cloze items (Szewczyk and Federmeier, 2022; Shain et al., 2024). Indeed, several studies have reported that LLM-derived predictions correlate strongly with N400 data (Heilbron et al., 2022; Michaelov et al., 2024), and some even suggesting that LLMs may outperform humans (Michaelov et al., 2022), especially when obtaining probability estimates for low-cloze items.

Building on our results that semantic cloze better predicts N400 amplitude than traditional lexical cloze, we next evaluated whether LLM-derived probabilities offer any advantage in modeling N400 responses. If the N400 truly reflects primarily semantic preactivation, we would predict that LLM-based lexical probabilities would not outperform semantic cloze in explaining N400 amplitudes.

## 3 Comparison with LLM-generated Probabilistic Measures

### 3.1 Rationale and Overview

As discussed in Section 2.6, recent work suggests that large language models (LLMs) can provide more accurate estimates of predictability than human-generated cloze norms (Michaelov et al., 2022), particularly for items with low cloze probability (Szewczyk and Federmeier, 2022; Shain et al., 2024). While direct comparisons between human- and LLM-generated cloze estimates in predicting N400 data remain relatively scarce, several recent studies have employed LLM-derived probabilities in place of traditional cloze tasks when predicting N400 amplitude (e.g., Heilbron et al., 2022; Szewczyk and Federmeier, 2022).

In principle, the relatively poor performance of lexical cloze as a predictor variable in our comparisons might be explained by human participants’ failure to capture the full range of low-cloze continuations. To investigate whether LLM-based probabilities can outperform our proposed semantic cloze measure, we compared both human- and model-derived estimates. Specifically, we focused on three LLMs that have been reported to exceed human predictability judgments — GPT, RoBERTa, and ALBERT (Michaelov et al., 2022).

Because LLMs compute probability distributions over tokens—a unit of text that may be a full word, subword, or special character—their predictions remain inherently lexical or sublexical in nature. While autoregressive models like GPT-2 estimate probabilities for the next token, bidirectional models such as BERT, ALBERT, and RoBERTa predict masked tokens within a sequence, but still at the token level. In other words, LLM-derived probabilities essentially represent a data-driven estimation of lexical cloze. Therefore, if the N400 primarily indexes semantic preactivation, we would not expect LLM-based probabilities to surpass semantic cloze in predicting N400 amplitude. To evaluate the performance of LLMs, we repeated our AIC analysis, now adding models with LLM-generated probabilities as predictors for comparison.

### 3.2 Computing LLM Cloze Probability & Surprisal

We calculated cloze probabilities for each target word in the ARK and NI18 stimulus sets using four large language models: GPT-2 (Radford et al., 2019), GPT-2.7b (Black et al., 2021), ALBERT (Lan et al., 2020), and RoBERTa (Liu et al., 2019). Although we were unable to obtain GPT-3 estimates for a direct comparison to (Michaelov et al., 2022) due to OpenAI licensing requirements, GPT-2.7b offers a reasonable proxy. It shares a similar transformer-based architecture and scaling principles, having been developed as an open-source model to replicate aspects of GPT-3’s performance (Black et al., 2021).

#### GPT-2 and GPT-2.7b

GPT-2 and GPT-2.7b (Radford et al., 2019; Black et al., 2021) are autoregressive language models that generate text by predicting the next token based on preceding context, leveraging a deep transformer architecture (Vaswani et al., 2023). Sentences were tokenized with the GPT-2 tokenizer, then truncated at the target word. We generated a probability distribution over possible next tokens based on the preceding context. In cases where target words were split into several tokens, probabilities were calculated iteratively: each sub-token’s probability was derived as a function of its preceding context, and the total probability was obtained by aggregating the individual probability values of all sub-tokens. This approach ensured accurate and context-sensitive measurement of surprisal values for both single-token and multi-token targets.

#### ALBERT and RoBERTa

ALBERT and RoBERTa (Lan et al., 2020; Liu et al., 2019) are bidirectional masked language models of the BERT family. Each full sentence was tokenized while masking the target region. The process replaced the target word with a mask token, and the probability distribution over vocabulary tokens for the masked position was computed. The probabilities of the original target word’s tokens were then extracted from this distribution. As with GPT, the probabilities of sub-tokens were aggregated when a target word was split.

All computations were implemented using the Hugging Face Transformers library Wolf (2020). The models were accessed via pre-trained weights, and tokenization was handled with the corresponding tokenizer classes. Computations were performed on GPUs for efficiency, leveraging PyTorch for all operations.

### 3.3 Analysis

We employed the same linear mixed-effects modeling framework described earlier, fitting separate models for each LLM-based predictor (GPT-2, GPT-2.7b, ALBERT, RoBERTa). Following that, we compare all models to each other and to the lexical and semantic cloze models using AICc/AIC.

Following common practice in research using LLMs, we also converted probabilities to word surprisal (*−log*_2_*P* (*Word|Context*) for both LLM-generated and human derived (lexical and semantic) cloze values). We present analysis for predictors expressed both as cloze probability and word surprisal.

Because zero cannot be log-transformed, we replaced zero-cloze values in human-generated measures with half of the lowest non-zero cloze probability observed in their respective dataset (0.0185/2 for ARK, 0.016/2 for NI18). This method provided a minimal but non-zero probability that could be log-transformed. All resulting surprisal values were then z-transformed and centered prior to analysis. Finally, we back-transformed the *β*-coefficients and 95% confidence intervals (CIs) from the z-scored scale into “raw” voltage units, reflecting the change in N400 amplitude associated with a hypothetical shift from 0% to 100% cloze probability.

### 3.4 Results

#### ARK

Tables 4 and 5 summarise the output of linear mixed effects modeling and AICc comparisons, for cloze probabilities and surprisal, respectively.

**Table 4.**
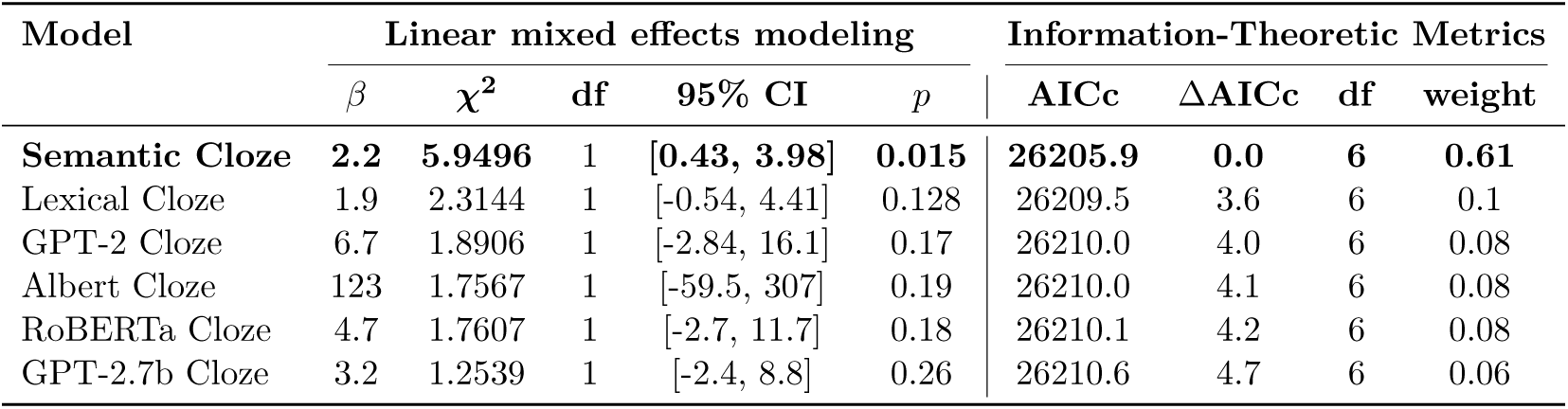
Output of linear mixed effects model comparisons and AICc. The models are ordered by their ΔAICc values from lowest to highest.

**Table 5.**
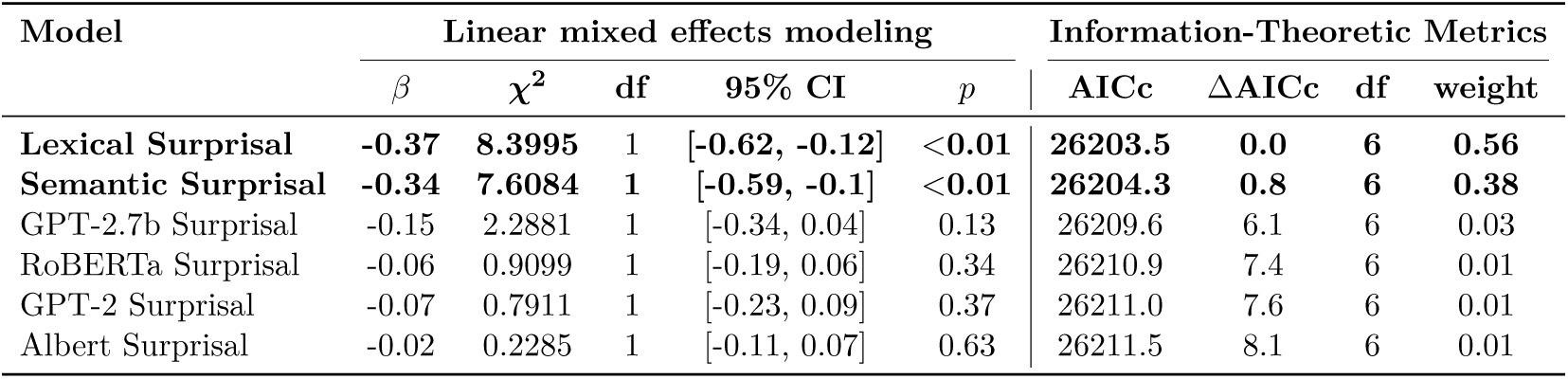
Output of linear mixed effects model comparisons and AICc. The models are ordered by their ΔAICc values from lowest to highest.

As in our previous analysis, semantic cloze emerged as the strongest predictor of the N400, as indicated by its minimal ΔAICc value. In contrast, the ΔAICc values for the GPT-generated cloze models (4.0 and 4.1) were comparable to that for lexical cloze (3.4), suggesting that these models provided similar predictive performance for N400 amplitude. Overall, however, all LLM-based and lexical cloze models received considerably less support than semantic cloze.

When predictors were expressed as surprisal (i.e., using the negative log transformation), the lexical surprisal model emerged as the best-fitting model—although it was only marginally better than the semantic surprisal model (ΔAICc = 0.8). Such a small ΔAICc value over the next-best predictor (GPT-2.7b) indicates that both models have substantial support in explaining the N400 data. Notably, the negative log transformation rendered the lexical surprisal model a significant predictor. In contrast, the transformation adversely affected the LLM-generated predictors: GPT-2.7b and Albert showed considerably less support, and GPT-2 and Roberta essentially no support according to the Burnham and Anderson heuristic.

#### NI18

Tables 6 and 7 summarize the linear mixed-effects modeling outcomes and AIC comparisons for cloze probabilities and surprisal values, respectively.

**Table 6.** Output of linear mixed effects model comparisons and AIC. The models are ordered by their ΔAIC values from lowest to highest.

| Model | Linear mixed effects modeling |  |  |  |  | Information-Theoretic Metrics |  |  |  |
| --- | --- | --- | --- | --- | --- | --- | --- | --- | --- |
| | $\beta$ | $\chi^2$ | df | 95% CI | $p$ | AIC | $\Delta\text{AIC}$ | df | weight |
| Semantic Cloze | 2.8 | 306.14 | 1 | [2.45, 3.07] | <0.001 | 192041.7 | 0.0 | 13 | 1 |
| Lexical Cloze | 2.7 | 287.24 | 1 | [2.35, 2.96] | <0.001 | 192060.5 | 18.8 | 13 | <0.001 |
| RoBERTa Cloze | 2.8 | 188.59 | 1 | [2.40, 3.19] | <0.001 | 192158.9 | 117.2 | 13 | <0.001 |
| GPT-2.7b Cloze | 2.7 | 167.36 | 1 | [2.3, 3.1] | <0.001 | 192180.2 | 138.5 | 13 | <0.001 |
| GPT-2 Cloze | 3.8 | 115.43 | 1 | [3.08, 4.44] | <0.001 | 192231.9 | 190.2 | 13 | <0.001 |
| Albert Cloze | 3.7 | 7.0465 | 1 | [0.97, 6.37] | <0.001 | 192340.3 | 298.6 | 13 | <0.001 |

Unlike in the ARK dataset, all models significantly predicted N400 amplitude in NI18, with two LLMs (RoBERTa, GPT-2.7b) yielding effect-size estimates (*β* = 2.8 and 2.7, respectively) comparable to human-based metrics.

Nevertheless, AIC comparisons indicate that none of the LLM-based models outperformed semantic or lexical cloze. In the cloze analyses (Table 6), RoBERTa was the next-best LLM with a ΔAIC of 117.2, and in the surprisal analyses, GPT-2.7b was the next-best with a ΔAIC of 45.1 (Table 7)—both exceeding 10 by a large margin. Consequently, these models have essentially no support relative to the top two predictors—semantic and lexical cloze.

**Table 7.**
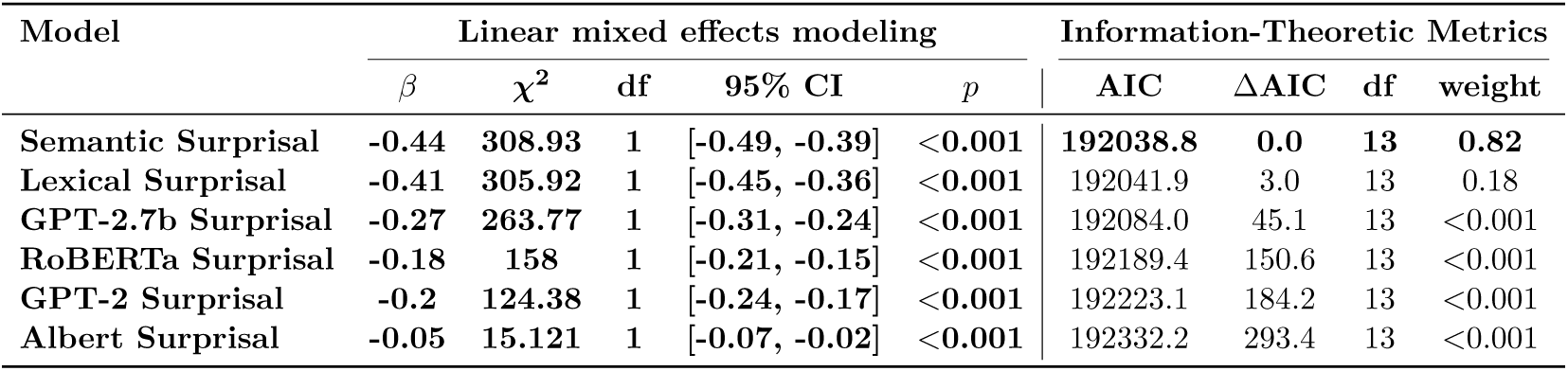
Output of linear mixed effects model comparisons and AIC. The models are ordered by their ΔAIC values from lowest to highest.

| Model | Linear mixed effects modeling |  |  |  |  | Information-Theoretic Metrics |  |  |  |
| --- | --- | --- | --- | --- | --- | --- | --- | --- | --- |
| | $\beta$ | $\chi^2$ | df | 95% CI | $p$ | AIC | $\Delta\text{AIC}$ | df | weight |
| Semantic Surprisal | -0.44 | 308.93 | 1 | [-0.49, -0.39] | <0.001 | 192038.8 | 0.0 | 13 | 0.82 |
| Lexical Surprisal | -0.41 | 305.92 | 1 | [-0.45, -0.36] | <0.001 | 192041.9 | 3.0 | 13 | 0.18 |
| GPT-2.7b Surprisal | -0.27 | 263.77 | 1 | [-0.31, -0.24] | <0.001 | 192084.0 | 45.1 | 13 | <0.001 |
| RoBERTa Surprisal | -0.18 | 158 | 1 | [-0.21, -0.15] | <0.001 | 192189.4 | 150.6 | 13 | <0.001 |
| GPT-2 Surprisal | -0.2 | 124.38 | 1 | [-0.24, -0.17] | <0.001 | 192223.1 | 184.2 | 13 | <0.001 |
| Albert Surprisal | -0.05 | 15.121 | 1 | [-0.07, -0.02] | <0.001 | 192332.2 | 293.4 | 13 | <0.001 |

### 3.5 Discussion

In this final set of analyses, we compared LLM-derived cloze probabilities to both semantic and lexical cloze measures. Our goal was to assess whether improving the precision of lexical cloze (by generating it through large language models) would improve this variable’s fit to the N400 data. Overall, semantic cloze remained the strongest predictor, reinforcing the idea that the N400 primarily reflects semantic rather than strictly lexical predictability. We note, however, that in the ARK dataset semantic and lexical cloze performed similarly when log-transformed. The negative log transform redistributes probabilities, thereby diminishing some of the raw-scale differences that might otherwise favor semantic cloze. Despite this slight convergence under log-transformations, the broader pattern across both datasets still supports semantic cloze as the more robust predictor. Importantly, for this analysis, we do not find the evidence that LLMs improved the lexical cloze estimation.

Nevertheless, in the NI18 dataset, LLMs performed well enough to emerge as significant predictors, with two models producing effect sizes on par with human-based metrics. This finding is not surprising, given that the NI18 target word were concrete nouns, resulting in minimal differences between semantic and lexical cloze estimates. As anticipated, LLMs do well at estimating exact-word probabilities, which appears sufficient in contexts where most target words map to a single label, like in the NI18 dataset.

Notably, LLMs also fared worse than lexical cloze, the other human-derived metric, echoing previous results showing that human-generated estimates generally fit N400 data better than LLM-based measures (Szewczyk and Federmeier, 2022). On the other hand, our findings differ from those reported by (Michaelov et al., 2022), who found that GPT-3, RoBERTa, and BERT-derived surprisal outperformed human predictions on the same NI18 dataset. One potential reason is that Michaelov et al. excluded zero-cloze items from their analysis. In contrast, our analysis retained the full dataset. To rule out the possibility that our chosen N400 window influenced these outcomes, we ran an additional analysis using the original N400 window means from Michaelov et al. (see Tables S3 and S4 in the Supplementary Materials). The results did not change our conclusions, indicating that model performance remained consistent with the patterns reported here.

## 4 General Discussion

In this study, we examined whether semantic cloze — our proposed measure of semantic feature predictability—predicts N400 amplitude more effectively than either lexical cloze (exact-word probability) or LLM-derived probabilities. Across two independent datasets (ARK, NI18), encompassing nouns, verbs, and adjectives, semantic cloze emerged as the strongest predictor of N400 amplitude. This result suggests that comprehenders pre-activate semantic features rather than fully specified words during language processing, which is in line with models linking the N400 to semantic (Rabovsky et al., 2018; Brouwer et al., 2017, e.g.) rather than lexical prediction (Fitz and Chang, 2019, e.g.).

### Semantic Cloze as an Improved Measurement Instrument

One of the longest-standing assumptions in neurolinguistics is that predictability, operationalized as lexical cloze probability, is a key determinant of N400 amplitude. Since the 1980s, lexical cloze has been the standard measure for estimating how expected a word is in context in N400 studies, with little to no adjustment over decades of research. However, during this same period, theoretical accounts of the N400 have evolved significantly. Most contemporary models now converge on the idea that the N400 primarily reflects processes related to meaning, rather than specific word forms. This raises a fundamental inconsistency: if the N400 is primarily sensitive to meaning, why do we continue to use a measure that only quantifies the probability of encountering a specific lexical item? This theoretical misalignment suggests that lexical cloze has low measurement validity for capturing what the N400 actually reflects.

Measurement validity is crucial because imprecise predictors can introduce significant biases in statistical modeling. Lexical cloze, as a noisy and theoretically misaligned measure, likely introduces error that distorts effect size estimation. One key assumption of linear regression is that predictor variables should be measured without error. When predictors are imprecise, parameter estimates can become biased and inconsistent, leading to misleading conclusions. In particular, measurement error in predictors can inflate false positives (Brunner and Austin, 2014) or attenuate true effects, leading to misestimated effect sizes (Altman and Krzywinski, 2024). Our results illustrate just how substantial this issue can be: in the ARK dataset, lexical cloze was not even a significant predictor of N400 amplitude.

In contrast, semantic cloze may offer a theoretically grounded and empirically superior alternative. By capturing the semantic feature predictability of a word rather than its exact lexical form, semantic cloze aligns naturally with established findings in N400 research. For example, it can account for the semantic similarity effect (Federmeier and Kutas, 1999), where words that are semantically related but lexically distinct still modulate N400 amplitude. Lexical cloze, by definition, cannot capture such effects, as it only considers a single correct response. Similarly, semantic cloze explains why function words (e.g., “a”/“an”) show no or reduced N400 effects (Nieuwland et al., 2018; Nicenboim et al., 2020) (unless articles themselves express definiteness as a semantic feature as in Carter and Nieuwland, 2022): while their lexical predictability is high, they contribute little to semantic pre-activation. These examples highlight the broader validity of semantic cloze across a range of N400 phenomena where lexical cloze would fail entirely.

Semantic cloze has a further theoretical advantage for theories of predictive processing: it aligns with a more parsimonious account of predictive processing, under which instead of generating detailed hypotheses about all possible next words, the comprehender can pre-activate only the relevant upcoming semantic features. This conceptualisation of predictability sidesteps major criticism of prediction in language comprehension—that prediction is too computationally costly to be a plausible mechanism (Jackendoff, 2003; Huettig and Mani, 2016), as it would require comprehenders to pre-activate an overwhelming number of potential exact-word continuations. Thus, when conceived as semantic rather than lexical prea-ctivation, prediction becomes a feasible mechanism.

Given these findings, there is little justification for continuing to rely on lexical cloze as the default measure of predictability in N400 research. Our results across two independent datasets suggest that semantic cloze may be a stronger, more precise predictor, and its use should be encouraged in future work. Of course, further studies should be done to validate these findings on additional datasets, but the fact that our results hold across two datasets already represents a substantial improvement over prior approaches. If the field aims to improve the precision and theoretical validity of its measurement tools, shifting toward semantic cloze as the default measure of predictability represents an essential step forward.

Our implementation of semantic cloze should be regarded as a proof of concept. Estimating semantic predictability via two human raters is admittedly labor-intensive. Future work should therefore explore more automated methods—ideally those that capture both the semantic predictability of content words and the lack of semantic predictability for function words (e.g., articles)—to refine the approach and further expand the utility of semantic cloze in N400 studies.

### LLMs and Cloze Probability

Our final analysis using large language models did not find that they performed better than semantic cloze. This is not surprising, given that LLMs operate at the level of word forms or sub-word token. By design, LLMs compute lexical probabilities, whereas the N400 appears to primarily reflect semantic processing.

This is not to say LLMs capture no semantic information. Research on BERT-like architectures, for instance, shows that hidden layers can encode syntax, semantics, and even real-world knowledge (Rogers et al., 2021). This information is distributed across different components of the model, from embeddings to attention heads, and manifests to different extents across layers. Alternatively, others (Slaats and Martin, 2025) demonstrate that information beyond word-form (i.e. syntactic, in their case) is able to indirectly influence model estimates without those features being necessarily encoded. In sum, whether the model’s hidden layers do encode some semantic or syntactic knowledge—or whether they merely integrate it indirectly—such information undeniably influences the final lexical distribution.

However, crucially, the final predictability estimate is drawn from a form-based distribution, assigning probabilities to specific word forms rather than partial semantic matches. Hence, while LLM-based estimates can approximate semantic probabilities in certain contexts, they do not generally outperform a semantic measure like semantic cloze for predicting N400 amplitude.

Although this lexical orientation limits their ability to capture N400 responses in contexts with multiple near-synonyms or abstract features, LLMs can still be valuable predictors in certain settings. In the NI18 dataset—which contained only concrete nouns as target words—LLM-derived probabilities correlated significantly with the N400, in some instances matching human-based effect sizes. When a target word maps to a single, canonical label (e.g., “apple”, “kite”), the distinction between semantic and lexical cloze narrows, allowing LLMs to perform relatively well.

However, for studies involving more semantically diverse or synonymous targets such as the ARK dataset, LMMs evidently perform rather poorly. Overall, caution is warranted before substituting human cloze norms with LLM estimates.

It is not our intention to discourage the use of LLM-generated predictability measures entirely. Rather, we suggest that future work might focus on refining LLM measurement to extract semantic cloze estimates. For instance, LLM word embeddings appear to capture contextually-dependent semantic information (Goldstein et al., 2022). By moving beyond the strict token-level probabilities, it may be possible to approximate or even replicate the advantages we observe with semantic cloze.

## Conclusion

Taken together, our findings demonstrate that semantic cloze provides a more precise and theoretically valid estimate of the predictability that drives N400 amplitude than either traditional lexical cloze or LLM-based cloze measures. Across two independent datasets (ARK and NI18), semantic cloze more consistently aligned with the N400’s apparent sensitivity to meaning, rather than to the form-based probabilities that LLMs and standard lexical cloze capture.

We have argued that in the context of N400 studies predictability should be reconceptualized in terms of semantic features rather than specific lexical items. Future research could explore methods for automating the estimation of semantic cloze—potentially leveraging large language models to capture semantic rather than merely lexical predictability. It would also be valuable to investigate whether semantic predictability exerts effects on behavioural measures such as eye-movements or reaction times in reading. Finally, we hope that other researchers will attempt to replicate these findings with different materials to independently evaluate the practical utility and robustness of semantic cloze. By moving away from purely lexical metrics, N400 studies can more accurately assess the role of meaning in language processing.

## Supplementary Material: Stimuli

**Table S1.** Example stimuli from the ARK dataset. The target word is in bold. The first two context sentences were presented together on the screen until button press. Button press initiated word-by-word presentation of the last sentence. There were no commas around ‘nonetheless’ to avoid effects of prosody.

| Distance | Predictability | Example Sentence |
| --- | --- | --- |
| Long | Expected | Peter came home from work with a terrible headache. He took some paracetamol before going to bed. Nonetheless his headache **persisted**. |
| Long | Unexpected | Peter came home from work with a terrible headache. He couldn't find any paracetamol at home. Nonetheless his headache **persisted**. |
| Short | Expected | Peter came home from work with a terrible headache. He took some paracetamol before going to bed. His headache nonetheless **persisted**. |
| Short | unexpected | Peter came home from work with a terrible headache. He couldn't find any paracetamol at home. His headache nonetheless **persisted**. |

**Table S2.** Example stimuli from the NI18 dataset. The target word is in bold.

| Condition | Example Sentence |
| --- | --- |
| Unexpected | The old wives’ tale says that if you want to keep the doctor away then you should eat a **carrot**. |
| Unexpected | For the snowman’s eyes, the children used two pieces of coal, and for its nose they used an **apple**. |
| Expected | The old wives’ tale says that if you want to keep the doctor away then you should eat an **apple**. |
| Expected | For the snowman’s eyes, the children used two pieces of coal, and for its nose they used a **carrot**. |

## Supplementary Material: Original NI18 Data Analysis

Here we repeat out analysis using the original N400 means provided by (Nieuwland et al., 2018), which are publicly available at https://osf.io/eyzaq/files/osfstorage.

**Table S3.** Output of cloze probability linear mixed effects model comparisons and AIC for NI18 using the original NI18 N400 means. The models are ordered by their ΔAIC values from lowest to highest.

| Model | Linear mixed effects modeling |  |  |  |  | Information-Theoretic Metrics |  |  |  |
| --- | --- | --- | --- | --- | --- | --- | --- | --- | --- |
| | $\beta$ | $\chi^2$ | df | 95% CI | $p$ | AIC | ΔAIC | df | weight |
| Semantic Cloze | 2.3 | 242.71 | 1 | [1.98, 2.55] | <0.001 | 187965.9 | 0.0 | 13 | 1 |
| Lexical Cloze | 2.2 | 226.15 | 1 | [1.89, 2.46] | <0.001 | 187982.4 | 16.5 | 13 | <0.001 |
| GPT-2.7b Cloze | 2.1 | 118.2 | 1 | [1.72, 2.47] | <0.001 | 188090.1 | 124.2 | 13 | <0.001 |
| RoBERTa Cloze | 2 | 116.68 | 1 | [1.68, 2.42] | <0.001 | 188091.6 | 125.7 | 13 | <0.001 |
| GPT-2 Cloze | 2.7 | 70.863 | 1 | [2.1, 3.35] | <0.001 | 188137.4 | 171.5 | 13 | <0.001 |
| Albert Cloze | 2.6 | 7.0465 | 1 | [0.14, 5.07] | <0.001 | 188203.9 | 238.0 | 13 | <0.001 |

**Table S4.**
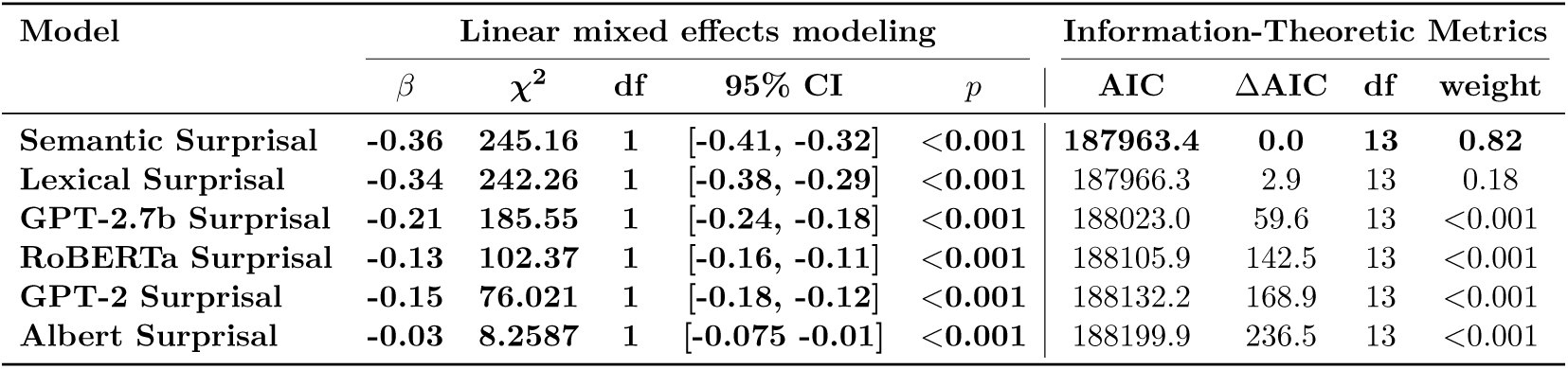
Output of surprisal linear mixed effects model comparisons and AIC for NI18 using the original NI18 N400 means. The models are ordered by their ΔAIC values from lowest to highest.

## Supplementary Material: List of NI18 changes

**Table S5.** Changes to the NI18 data introduced in the present study.

| Parameter | Nieuwland et al. | This manuscript | Comments |
| --- | --- | --- | --- |
| Lexical cloze probability for item 79 (exp) | 97% | 100% | In NI18 one cloze response was missing, yet cloze was still computed over 30 participants. We removed the missing response and recalculated over 29 participants. |
| Baseline correction | −100–0 ms | −200–0 ms | NI18 used a 100 ms baseline to mirror DeLong et al. (2005). We opted for the more traditional 200 ms baseline, which also matches the preprocessing for ARK. |
| N400 window | 200–500 ms | 350–450 ms | NI18’s window replicated DeLong et al. The 350–450 ms interval is the recommended N400 window and ensures consistency with ARK. |
| Number of datapoints | 25 848 | 25 846 | Two trials contained mis-timed EEG triggers (likely hardware glitches). Because the triggers did not reliably mark word onset, we excluded these trials from the final data set. |

## Footnotes

1 The choice of N400 window and electrodes in the original (Nieuwland et al., 2018) study was motivated by replicating parameters from DeLong et al. (2005). NI18 extracted N400 means from the 200–500 ms window over electrodes Cz, C3/4, Pz, and P3/4. Here, we used their pre-processed EEG data to extract new means from the 300–500 ms window to maintain consistency across both datasets. The 300–500 ms window is the recommended N400 measurement window (Kappenman et al., 2021) for this component. To make sure our results were not driven by our choices of the N400 window and electrodes, we have additionally repeaded our analyses on the N400 means provided by the original authors (see Supplementary Materials).

## Notes

https://osf.io/eyzaq/files/osfstorage

### Competing Interest Statement

The authors have declared no competing interest.

### Summary of Updates

Fixed a reference syntax; Wording about 'explained variance' changed to 'model fit' in the abstract and the results section

### Funding

- Deutsche Forschungsgemeinschaft (DFG, German Re-search Foundation) – Project ID 317633480 (SFB1287)

